# Cell delivery peptides for small interfering RNAs targeting SARS-CoV-2 new variants through a bioinformatics and deep learning design

**DOI:** 10.1101/2022.02.09.479755

**Authors:** Ricardo D. González, Pedro R. Figueiredo, Alexandra T. P. Carvalho

## Abstract

Nucleic acid technologies with designed delivery systems have surged as one the most promising therapies of the future, due to their contribution in combating SARS-CoV-2 severe disease. Nevertheless, the emergence of new variants of concern still represents a real threat in the years to come. It is here that the use of small interfering RNA sequences to inhibit gene expression and, thus, protein synthesis, may complement the already developed vaccines, with faster design and production. Here, we have designed new sequences targeting COVID-19 variants and other related viral diseases through bioinformatics, while also addressing the limited number of delivery peptides by a deep learning approach. Two sequences databases were produced, from which 62 were able to target the virus mRNA, and ten displayed properties present in delivery peptides, which we compared to the broad use TAT delivery peptide.

## INTRODUCTION

The current severe acute respiratory syndrome coronavirus type 2 (SARS-CoV-2) is a virus that caused the 2019 outbreak of coronavirus disease (COVID-19) pandemic, with origin in Wuhan, China [1]. The success of the new rapidly developed vaccines has propelled the interest in nucleic acids-based therapies as an approach to treat several systemic disorders, with an underlined interest in ribonucleic acids (RNA) [1]. However, new variants of concern have emerged, such as B.1.617.2 (Delta) and, more recently, the B.1.1.529 (Omicron) variants, which have become dominant in an international setting, with higher transmissibility rates and consequently higher affluence of patients to health systems [2]. Given that, new approaches to reduce viral charge in infected individuals are needed to diminish infection rates and, possibly, symptoms. In that concern, small interfering RNA (siRNA) have been extensively studied and are becoming an accepted modality of pharmacotherapy [3]. These short strands of RNA induce selective gene suppression by binding to sequence-specific messenger RNAs (mRNA) leading to their cleavage and degradation, thus preventing protein synthesis [4].

Nevertheless, the delivery of nucleic acids imposes challenges like efficient cell delivery. New delivery methods focus on the design of nanoparticles that will protect the RNA from degradation and allowing the crossing of the cell membranes for intracellular delivery, such as lipid nanoparticles, polymer-based nanocarriers (nanoparticles, micelles-based) and carbon-based nanostructures [5]. Nevertheless, protein transduction domains, most commonly known as cell penetrating peptides (CPP), are small peptides (6 to around 30 amino acids) that are able to carry cargos across the cellular membranes in an intact and functional fashion [6]. They have been used complementary to the aforementioned methods, allowing an enhanced endosomal escape by conjugation on the surface of those structures (nanoparticles, polymeric micelles, or liposomes). These complex constructs need a thorough design and assembly, which may hinder the development of stable RNA therapies and delivery.

Nucleic acids have been successfully delivered to cells by CPPs [7], but how this occurs remain unclear among researchers. They are mainly cationic in nature but can also have amphiphilic properties [8], though no clear rules to distinguish if a peptide is a CPP or not are clearly established, limiting the number of described peptides. Even for the same CPP, the transduction mechanism varies depending on the cargo, cell environment and peptide concentration. Given that, CPPs have been described to be mostly internalized by endocytosis even if different results have made it difficult to indicate with precision which endocytic pathway is involved [9,10]. The most used CPPs: trans-activator of transcription (TAT) protein of HIV-1, penetratin and transportan [9], have been used in preclinical and clinical studies on Alzheimer’s disease, cancer, or cerebral ischemia for the delivery of therapeutics, but caution due to cytotoxic effects must be ensured [11]. Notwithstanding, even if their mechanism is not clear, CPPs seem to be a great approach for the delivery of therapeutics, namely siRNAs, due to their smaller size [3].

Here, we have designed siRNA sequences for COVID-19, considering the new variants of concern that have been spreading worldwide. We have also addressed the limited number of CPPs by applying a deep learning approach for the design of new peptides capable of delivering siRNAs to inhibit RNA activity, thus reducing viral replication. Moreover, insights in the absorption, distribution, metabolism, and excretion (ADME)/toxicity of the peptides were accessed, comparatively to the most used TAT delivery peptide.

## METHODS

### siRNA sequence library

An siRNA sequence library was constructed from the the SARS-CoV-2 reference genome (NCBI, NC_045512), based on the protocol proposed by Medeiros *et al.* [12]. A 21-array sliding window approach with steps of 1 was implemented to obtain sequences with 21 nucleotides (nt). These sense sequences were then transcribed to their antisense counterpart via shell scripting to allow proper targeting, followed by filtering steps. As our purpose was to obtain sequences with siRNA features, a proper selection was necessary. Biological activity was controlled by restricting GC-content to 30-50%, and by removing sequences able to form hairpins and of self-annealing, whereas toxicity was evaluated by analyzing the presence of poly(U/T/A) and GCCA motifs [13]. We wanted these sequences to be able to target SARS-CoV-2 and its Delta and Omicron variants, but also to other related viral diseases, such as SARS-CoV, Middle East Respiratory Syndrome-related coronavirus (MERS-CoV), and influenza (H1N1), with no off-target effects in humans. To do that, we retrieved the following genomes in FASTA format: 1) the human genome, coding and non-coding transcriptome (GRCh38, from NCBI and ENSEMBL, respectively); 2) the reference genomes of SARS-CoV-2 (NC_045512.2), SARS-CoV (NC_004718.3), MERS-CoV (NC_038294.1), and influenza (GCF_001343785.1); and 3) different SARS-CoV-2 variants (original 2019 variant and the currently dominant Delta and Omicron) from USA (Texas, California and New York), Brazil, Portugal, Spain, England, Germany, Russia, China (without Wuhan), and Wuhan strains, obtained from the Global Initiative on Sharing Avian Influenza Data (GISAID) [14]. Alignment to those genomes was performed using the short-reads aligner Bowtie v1.1.0 [15], reporting all valid alignments per read, with the following parameters: maximum number of attempts to match an alignment = 4, and maximum number of mismatches in the “seed” = 3, with “seed” length = 7. The analysis of these features allowed for the selection of the most promising sequences capable of targeting different genes of interest in COVID-19, acting as siRNAs, while being able to target related viral diseases, enhancing the targeting scope of these sequences.

### Delivery peptides design

The design of CPPs was performed according to the protocol on antimicrobial peptides described by Tucs *et al.* [16]. This method relies on deep learning generative adversarial networks (GAN), which control the probability distribution of the newly generated sequences, ranking them into positive and negative classes. The underlined criteria include physicochemical descriptors such as charge, hydrophobicity, and molecular weight. Data was retrieved from the publicly available databases CPPsite 2.0 [17], a webserver with deposited and curated CPPs (data available as of 1^st^ October 2021), and machine learning CPP (MLCPP) [18], a two-layer prediction framework for machine-learning-based prediction of cell penetrating peptides. Additionally, as the work by Tucs and colleagues [16] on antimicrobial peptides showed that most of their new peptides were cationic and amphiphilic, which are features found on CPPs, and to enrich the training sets for GAN, sequences from their datasets (retrieved from APD, CAMP, LAMP and DBAASP databases) were also collected. Sequences with up to 52 amino acids were used for training. Redundant sequences were removed. The final dataset contained 14,778 positive sequences (CPP and antimicrobial peptides), and 6,664 negative sequences (non-CPPs). All the sequences were used during the training of the model (for activity prediction) followed by GAN of the positive dataset.

### ADME/Toxicity

Absorption, Distribution, Metabolism, Excretion and Toxicity (ADME-Tox) studies were conducted using the variable nearest neighbor (vNN) webserver for ADME prediction [19], which allows for the retrieval of a range of properties, such as cardio- and cytotoxicity, and the likelihood of causing liver injury. This *in silico* method permits for a first scan of compounds before taking new molecules to the lab, concomitantly having a great potential as its learning algorithms rely on available experimental data.

## RESULTS AND DISCUSSION

### Small interfering RNAs

Before the outcome of RNA vaccines, the regulatory mechanism achieved by RNA interference (RNAi) was already known and used for RNAi-based therapies [3]. These usually short RNA strands, such as siRNA, caused sequence-specific gene suppression with several advantages over other therapies: the sequence specificity allows for targeting of oncogenes and growth factors, and even to target single nucleotide polymorphisms [20]. Furthermore, upon proper delivery, siRNAs can be delivered to the brain, enhancing the scope of RNA interference to treat neurodegenerative pathologies, and to viral targets such as HIV and Hepatitis C, by inhibition of the viral RNA [21]. The vast potential of this technology to target viral diseases is of most importance in the current worldwide COVID-19 pandemic, and a great opportunity to further development of therapies for diseases that are based on differential gene expression. It is noteworthy that RNA technologies allow for a faster development compared to classical viral vaccines and are more easily modified to follow new variants.

The sliding window of one-step approach allowed to produce 29,880 21 nt-long sequences that could target some or most of the targeted genomes under study (available upon request). The alignment of those sequences to the genome of SARS-CoV-2 allowed for the annotation of 29,197 sequences to several genes of interest (Figure 2 and Supplementary Information Table S1).

**Figure 2.**
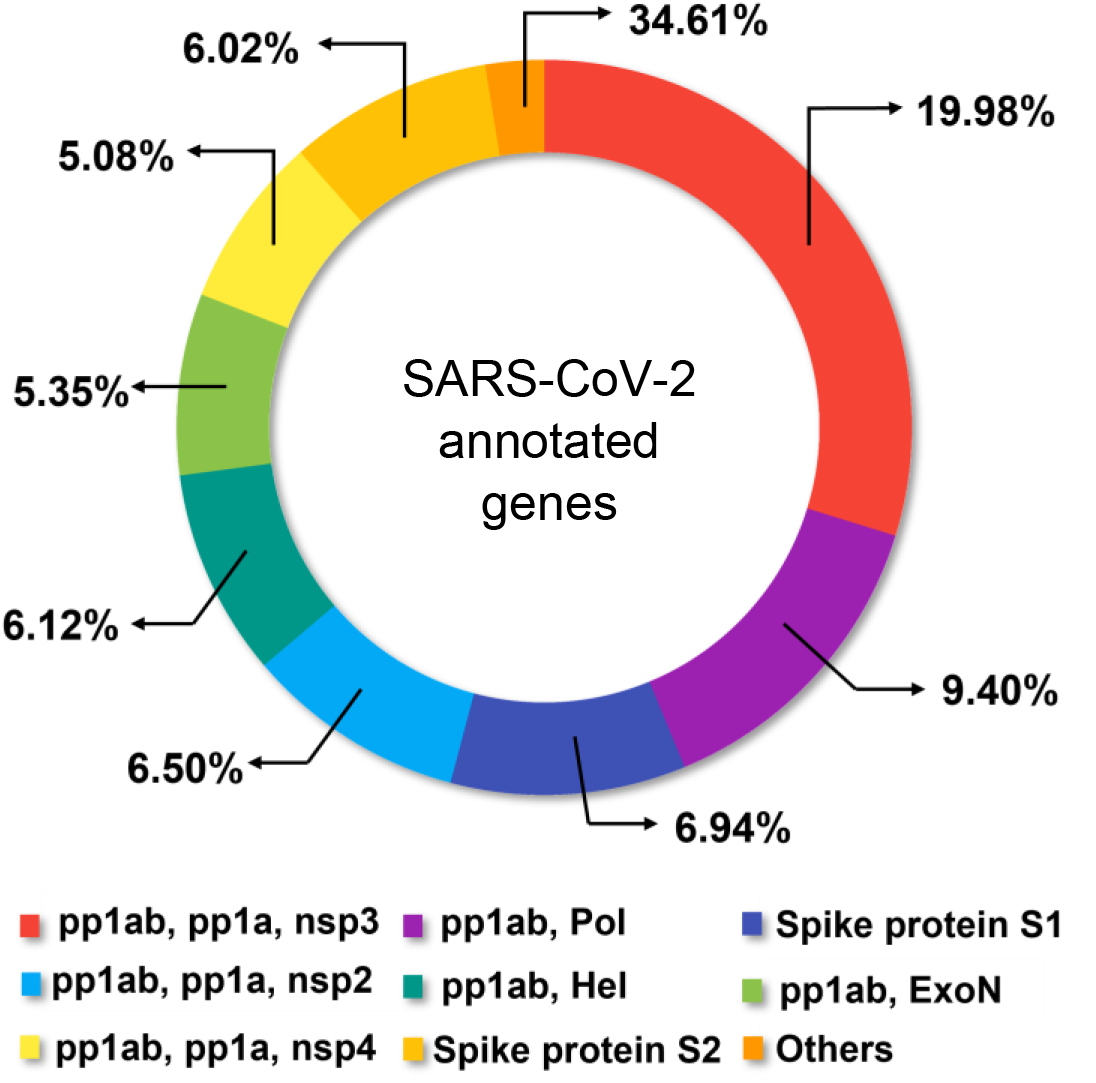
Distribution of annotated genes that the produced 21-nt sequences are able to target. Pp, polyprotein; nsp, non-structural protein; ExoN, exoribonuclease; Pol, polymerase; Hel, helicase. Genes with less than 3.5% representation were pooled in the “others” group and are described in Supplementary Information (Table S1).

SARS-CoV-2 genome is similar to other coronaviruses. It is a ssRNA that encodes 27 proteins from 14 open reading frames. Eight accessory proteins and the four main domains (nucleocapsid, envelope, membrane, and spike proteins) are encoded by the 3’-terminus, while polyproteins (pp) 1ab and 1a (upon cleavage of pp1ab) are encoded by the 5’-terminus. These are responsible for coding non-structural proteins (nsp) 1 to 10, whereas nsp11 is exclusively encoded by pp1a, and nsp13-16 from pp1ab [22].

Most of the produced siRNA sequences (≈ 20.0%) target the pp1ab gene and those it encodes (pp1a and nsp3). Other than spike proteins, most of the generated sequences target this 790 kDA polyprotein, as it is present in most gene sets (73.12%). Pp1ab and pp1a are replicases processed by two viral proteases (papain-like and 3C-like protease), important in viral replication and transcription, which makes them a vital target to inhibit viral activity [23]. On the other hand, only around 7.0% of the sequences aimed at the spike glycoprotein S1, and 6.0% to S2, totaling 13.0% of sequences to the spike protein. This protein is responsible for the binding to the host cell receptor and for the induction of membrane fusion, which made it a vital player in the host invasion process, mainly through the high affinity to the angiotensin converting enzyme (ACE) 2 receptor [24]. This is an interesting finding, as most current approaches to treat COVID-19 relied on targeting the latter, highly mutagenic protein, while it seems that pp1ab gene offers more targeting capability while ensuring the inhibition of the replication and transcription, without which no host invasion is possible.

To make sure that the designed siRNAs would have an activity close to those retrieved from experimental settings, we applied several filters to the obtained sequences. GC-content is an important factor to biological activity, and it has been found that a GC-content ranging from 30.0 to 50.0% enables better activity [25]. We have applied that scheme and came up with 19,731 sequences with GC-content of 33.0 to 48.0%, targeting the same annotated genes.

Next, poly(T/U) and poly(A) with more than four nucleotide repeats should be avoided, as these tags can act as termination signals to RNA polymerase III [26]. Thus, we excluded sequences with 4 or more of these repeats, ending up with 15,681 siRNA sequences.

Toxicity is also a major concern regarding new therapeutics, being them drug or nucleic acids-like therapeutics. According to Fedorov *et al.* [27], to limit toxicity, these sequences must have at least three mismatches compared to the human genome, and they should not possess poly(T/U) and GCCA tags (the first one has been resolved in the previous filtering step). We proceeded then to the removal of sequences with GCCA motifs and those that had less than three mismatches when aligned to the human genome and human coding and non-coding transcriptome. These filtering steps were performed by shell scripting and spreadsheets conditionals, and highly allowed for an 87.0% reduction on the number of siRNA sequences (1,993).

Apart from GC-content, effectivity was also assessed by removing 614 palindromic sequences, those capable of forming hairpin structures and of self-annealing, which would difficult or hinder the oligonucleotide’s binding to the RNA target. A final cleaning step was performed according to effectiveness prediction. To this aim, sequences were scanned in the si-shRNA selector program [28], after which only 68 siRNA sequences remained. This program identifies features associated with silencing efficiency, scoring the sequences through statistical structure-activity relationships based on thermodynamic parameters.

Finally, the 68 sequences respected all the filtering that we thought would make them more effective to our purpose. Nevertheless, we wanted those sequences to have a broad effect, by being capable to target all the strains of different countries and variants (original, Delta and Omicron), while also targeting other viral diseases related to SARS-CoV-2, such as SARS-CoV, MERS-CoV, and H1N1, which have similar genomes. Therefore, we looked at the number of alignments to each of these genomes against the siRNA sequences and removed those with no alignments at all. In total, out of the initial 29,880 sequences, we came up with 62 21-nt long sequences that could act as siRNA for those viral infectious diseases and variants (Supplementary Information, table S2). This last filtering removal was highly influenced by the number of matches to COVID-19 Omicron variant of concern, as there were not as much completed sequences with high coverage deposited in the GISAID platform, hence a total binding to the 21-nt sequences was lower. Despite the significant removal of the original siRNA sequences, the relative targeting to the genes of interest remains the same as in the original data set (around 20.0% for pp1ab, pp1a and nsp3 cluster, Figure 3). It is important to note that a general first step of siRNA sequences generation without filtering enables researchers to apply their own designed filters to select those sequences that would better fit their needs.

**Figure 3.**
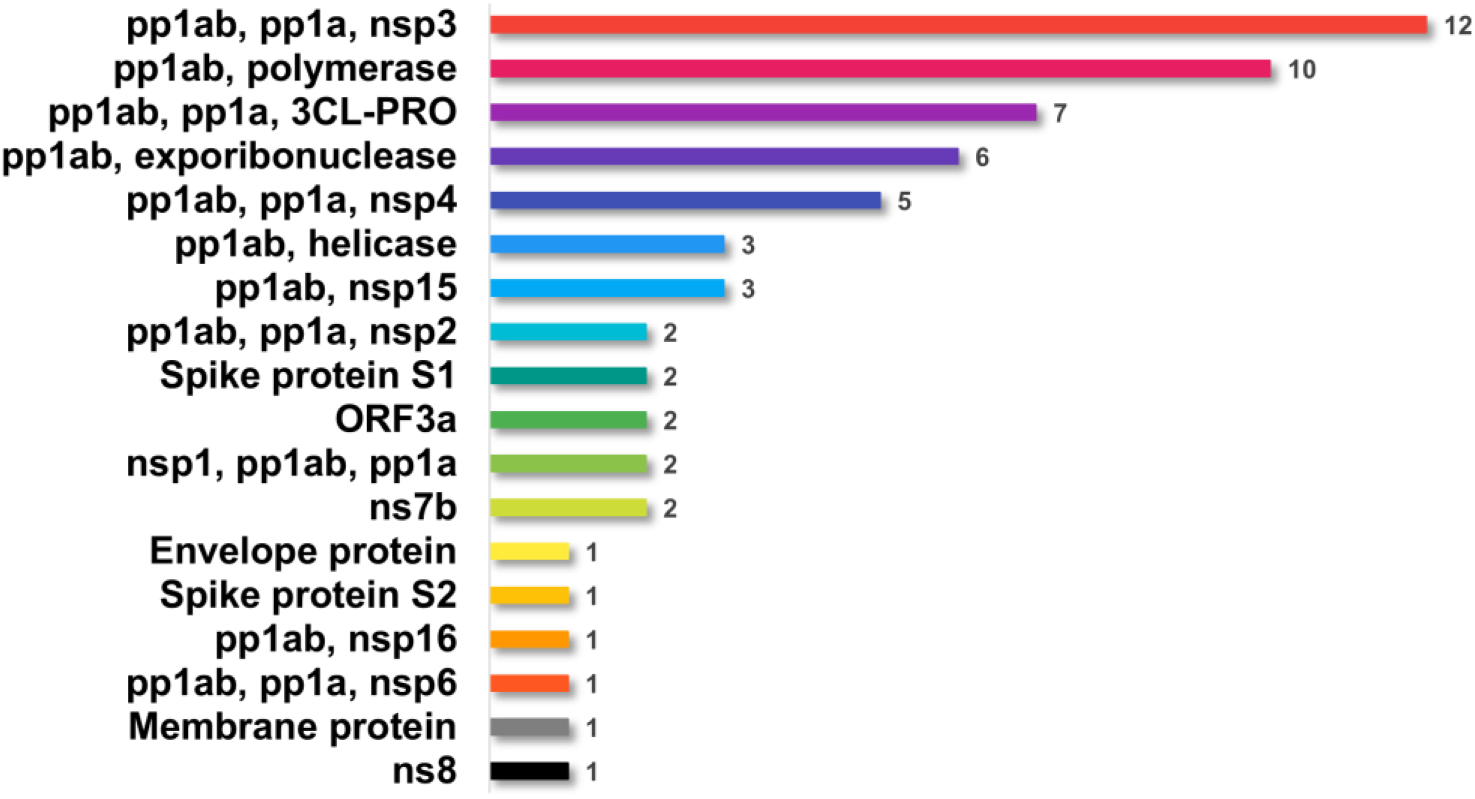
Distribution of annotated genes that the produced 21 -nt sequences are able to target, after screening of promising siRNA sequences. Pp, polyprotein; nsp, non-structural protein; 3CL-PRO, 3C-like protease; ORF, open reading frame; ns, accessory proteins.

### Cell Penetrating Peptides

The delivery of therapeutics to cells is a major focus in today’s therapy research, as to ensure the proper localized activity of those moieties without raising off-target effects. In that regard, several nanostructures have been suggested to protect the carried drug cargos; however, nucleic acids-like therapies focus on base pairing rather than structural features, such as those that are screened for drug repurposing. In that sense, it is possible to use simpler but effective transporters to deliver them. CPPs have the ability to cross membranes in a non-invasive manner, allowing for the maintenance of cells’ integrity while also being considered safe and highly efficient, showing low cytotoxicity and no immunological responses [8,11]. However, the number of described CPPs in the literature is scarce, and cell delivery is usually achieved with the broad CPPs TAT, penetratin and transportan [9]. Thus, we have here delved into deep learning methods to design new delivery peptides.

By controlling the probability distribution of the new peptide sequences, the deep learning GAN method allowed for the sorting of those sequences into positive (CPP) and negative (non-CPP) classes, considering physicochemical descriptors such as charge, hydrophobicity, and molecular weight of the data from which it learned.

New sequences were created, in a total of 9,984 (data available upon request), ranging from 5 to 52 aa. From those, we removed four sequences that contained the Asx (B) unnatural amino acid (either aspartic acid or asparagine were possible in those positions), totaling 9,980 new sequences. This step facilitated the use of the following servers, as they only accept natural amino acids sequences.

Despite the learning capability of these methods, we wanted to further filter our data. As a result, we have resorted to the server Machine Learning CPP (MLCPP), from which we initially extracted data for our learning datasets. This tool enables the discrimination of CPP and non-CPPs, scoring the sequences based on their amino acids’ composition. It classified 3,866 sequences as CPP and 6,114 as non-CPP. Alongside the probability score of being a CPP or not, which we used as classifier, the server also provides a probability score regarding uptake efficiency. Both are important parameters for the determination of effective delivery peptides, so we organized the sequences according to both conditions. Interestingly, at least up to the 15 best scored of each classifier, there was no sequences present in both. This highly suggests that being positively scored as a CPP, due to physicochemical properties, does not imply a high uptake efficiency score.

For further analysis, we have selected the top 5 sequences from each score classifier and performed a BLAST alignment to evaluate if these sequences were already deposited elsewhere in any genome. As no match was found, we assumed these sequences as new potential CPPs, and compared them to the results of the most used CPP, the TAT peptide, first by sequence alignment (Clustal Omega, Figure S1) [29] and by score analysis (Table 1).

**Table 1.**
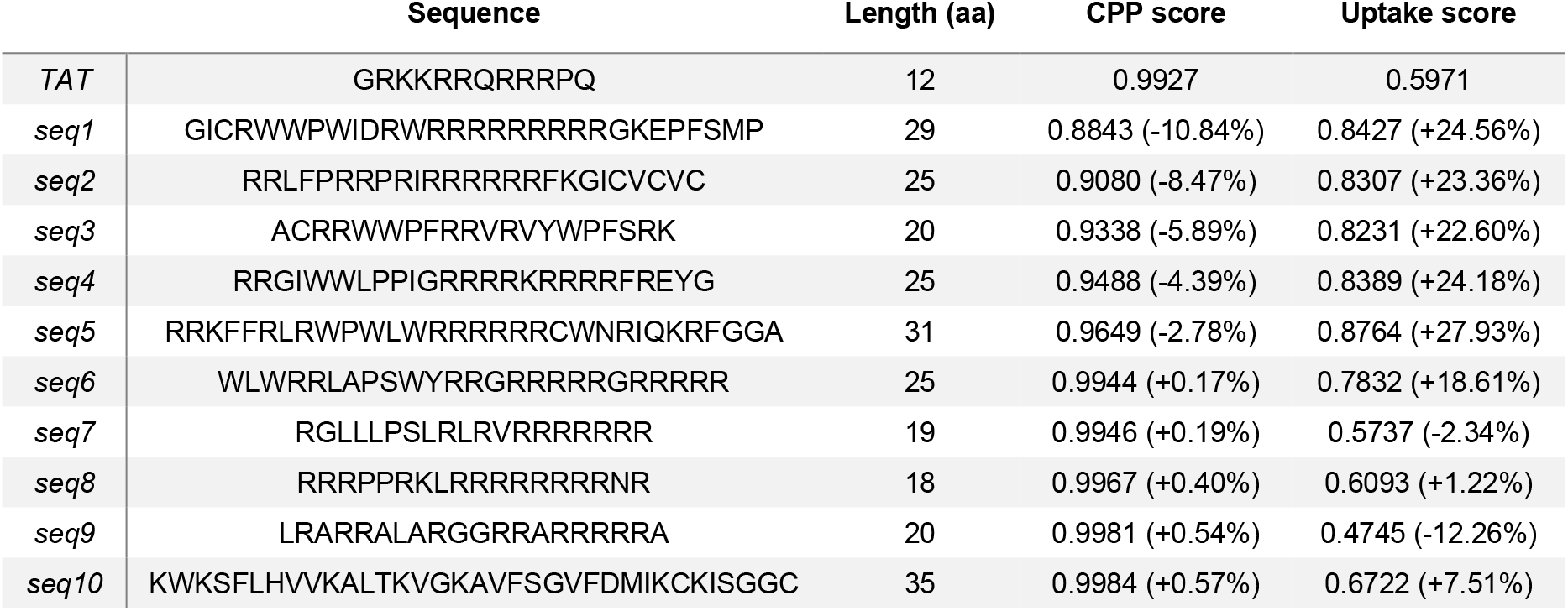
Uptake and CPP probability scores of the top 10 new sequences. Values in parenthesis concern to comparison to TAT.

Out of the 10 sequences, five displayed a slightly higher CPP score compared to TAT peptide (99.27% *versus* the higher 99.84%), while most of them scored higher than TAT regarding uptake efficiency, with only two sequences scoring lower (seq9 47.45% and seq7 57.37% *versus* TAT’s 59.71%). The sequence with highest uptake score (seq10) is up to 1.5-fold better than TAT. These scores, however, do not seem to be related with the length of the sequences, as no pattern could be found. They may rely, more than on sequence size, on amino acid composition and physicochemical features. The latter were then calculated using HeliQuest server [30] (Supplementary Information, Table S3), from which it was also possible to retrieve the peptides helical wheel representations (Figure 4).

**Figure 4.**
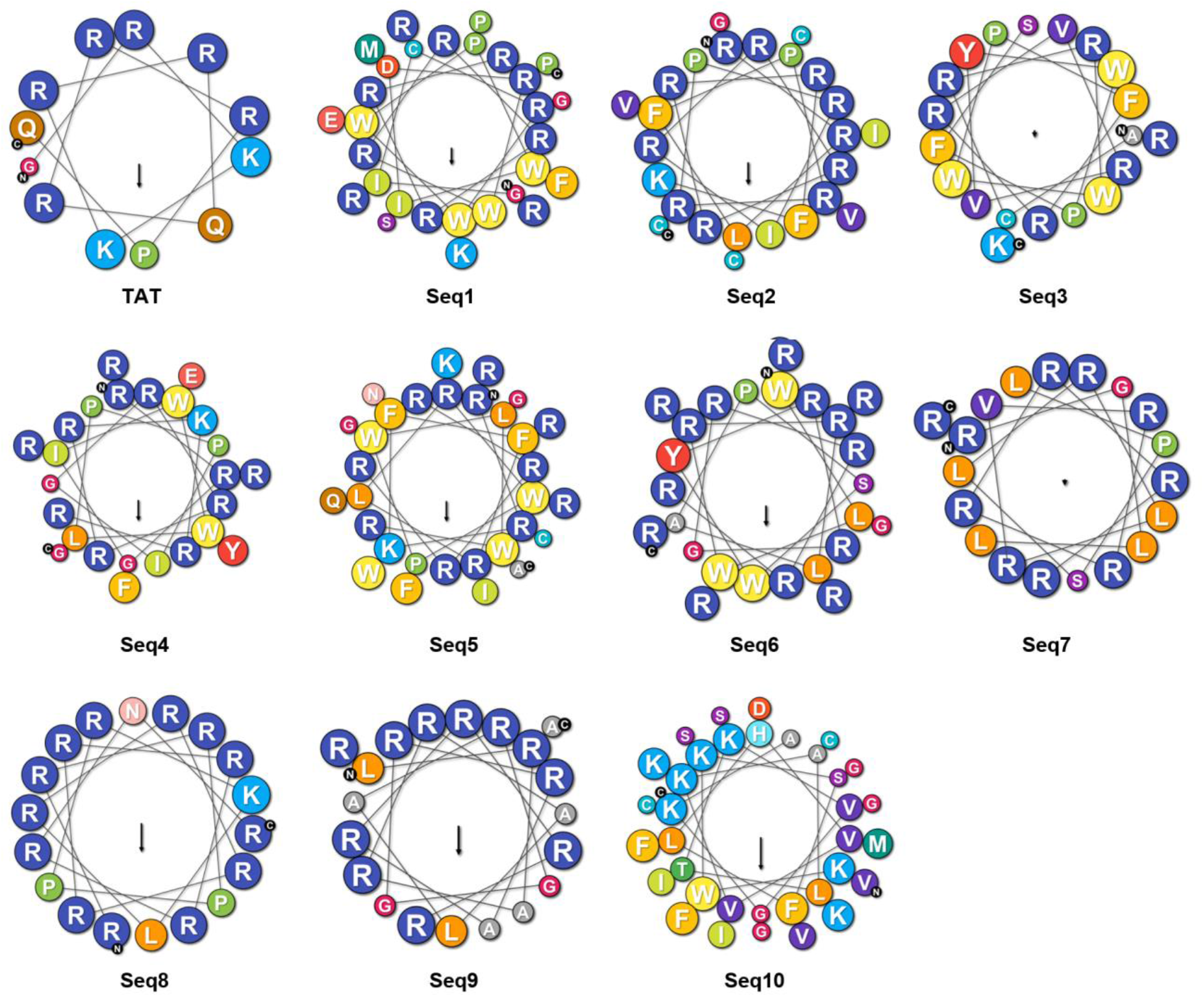
Helical wheel representation of the new peptides.

None of the sequences, including TAT, present highly amphipathic helices, with hydrophobicity inferior to 0.6. TAT is the peptide with higher polar residues content (91.67%), while the new peptides averaged at 61.07%. However, five of the new sequences (seq2, seq3, seq7, seq9, and seq10) have hydrophobic faces, which are said to be features of biologically active peptides, as these hydrophobic residues may insert between the membranes’ lipid acyl chains, allowing for binding and crossing the membrane [31].

### Citotoxicity

To obtain new effective and safe drugs to be used in clinical settings, the compounds need to be screened and characterized regarding their cytotoxicity. One of the most common studies performed to obtain this information is the ADME-Tox assay, which allows for the prediction of what will occur after administration in the human body. Despite being a highly *in vitro* process, several *in silico* pipelines have been created based on experimental results to more easily help researchers developing and characterize new products. Such an example is the variable nearest neighbor (vNN) webserver for ADME prediction [19], which allows for the retrieval of a range of properties, as cardiotoxicity, cytotoxicity, and the likelihood of causing liver injury. We have scanned our new sequences and the TAT control in the server, to observe how they compared to the latter, a broadly accepted delivery peptide (Table 2).

**Table 2.** ADME/Toxicity results for the top 10 new CPP sequences. MRTD, maximum recommended therapeutic dose (in mg/day, for an averaged adult weighting 60 kg). In parenthesis, fold-change compared to TAT.

|  | Cytotoxicity | Liver injury | Metabolism | CYP3A4 inhibitor | MRTD |
| --- | --- | --- | --- | --- | --- |
| TAT<sup>#</sup> | No | Yes | ≤ 30' | No | 16 |
| Seq1<sup>§,¶</sup> | No | No | ≥ 30' | Yes | 1445 (90.31) |
| seq2 | No | Yes | ≤ 30' | No | 1791 (111.94) |
| seq3<sup>§</sup> | No | No | ≥ 30' | Yes | 1112 (69.50) |
| seq4<sup>§</sup> | No | No | ≥ 30' | Yes | 1362 (85.13) |
| seq5 | No | No | ≥ 30' | Yes | 1747 (109.19) |
| seq6 | No | No | ≥ 30' | Yes | 1525 (95.31) |
| seq7 | No | No | ≤ 30' | No | 1817 (113.56) |
| seq8<sup>#</sup> | No | Yes | ≤ 30' | No | 391 (24.44) |
| seq9<sup>*</sup> | No | No | ≤ 30' | No | 2994 (187.13) |
| seq10<sup>§</sup> | No | No | ≥ 30' | No | 1596 (99.75) |

<sup>\*</sup> Cross blood-brain barrier; <sup>#</sup> P-glycoprotein inhibitor; <sup>§</sup> P-glycoprotein substrate; <sup>¶</sup> Ether-à-go-go-Related (hERG) gene (cardiotoxicity).

Regarding cytotoxicity, none of the new sequences were predicted to cause damage to cells, considering an IC_50_ ≤ 10.0 μM. However, three sequences, seq2, seq8 and TAT, have a high risk of inducing liver injury, which would withdraw the use of these peptides. These same peptides, plus seq9 and seq7, were predicted to be rapidly metabolized by the liver (half-life lower than 30 minutes). All the others, on the other hand, may be CYP3A4 inhibitors, which may justify their half-life being superior to 30 minutes, as they may be able to circumvent xenobiotics metabolism by this CYP enzyme. TAT and seq8 may be inhibitors of P-glycoprotein, while peptides seq1, seq4, seq3 and seq10 may be substrates. This protein is an essential membrane protein with the role of transporting foreign compounds from the cell to the outside. Given that, the latter set of peptides may not be ideal for cell delivery, as they could be extracted from inside the cell through the P-glycoprotein pathway – nonetheless being able to cross the cellular membranes. Interestingly, the TAT peptide has been assigned as capable of crossing the blood-brain barrier in several studies [32], but the server could not correctly predict this feature when stronger filters were applied, only when weaker filters were used. Furthermore, the peptide seq1 appears to have some cardiotoxicity (hERG). According to the server’s training data for this feature, this sequence must present an IC_50_ ≤ 10.0 μM. This peptide should then be avoided to prevent the blockage of a gene coding for a potassium ion channel involved in normal cardiac repolarization. None of the sequences showed mitochondrial toxicity or mutagenicity (through mitochondrial membrane potential and mutagenicity probability (Ames test), respectively). At last, it was important to verify the maximum recommended therapeutic dose (MRTD), this is, the daily dose that is considered safe to patients to take. Remarkably, TAT peptide has the smallest MRTD value of 16 mg/day, while seq9 has the highest MRTD of 2,994 mg/day, 188-fold better than the control.

This data suggests that seq7 and seq5 peptides may be the safest ones to cell delivery purposes. While the first may have a half-life prediction of up to 30 minutes, it respects all the other parameters, concomitantly being the second with highest MRTD (1,817 mg/day), which could make up for the shorter half-life. This peptide has a higher probability score of being a CPP (99.46% *versus* TAT’s 99.27%) and its probability score on uptake efficiency is close to that of TAT (57.37% *versus* TAT’s 59.71%). Seq5 also passes all the parameters except for the CYP3A4 inhibitor prediction. This parameter should be taken into account, as the inhibition of CYP3A4 activity may lead to increased concentration of the compound in the liver and the intestine, which could cause toxicity [33]. Nevertheless, no cito- or cardiotoxicity were predicted for this peptide, and it has the third higher MRTD value (1,747 mg/day), which could point to its general safety.

## CONCLUSIONS

The emergence of the viral COVID-19 disease has propelled the way to develop better and more personalized medicines, with the highlight to RNA therapeutics. This is a great time for delving into these new technologies and broad their scope to other pathologies.

Nevertheless, the design of therapeutics directed to nucleic acids sequences requires careful design in order to enhance their efficiency and also assure their safety and the lack of off-target effects that may interfere with other genes expression and, consequently, protein synthesis and function. In that sense, computational tools ranging from bioinformatics analysis to the more computationally expensive machine and deep learning have paved their way into the routine of researchers prior to the extensive and comprehensive experimental evaluations. Here, we have explored the genomics of SARS-CoV-2 to produce sequences able to bind to its genes to produce inhibitory effects, further preventing protein synthesis by the RNA interference pathway, leading to RNA degradation. The availability of data, in part caused by the huge demand of COVID-19 data, allowed us to define sequences with a broad general use. These new methodologies allow for the quick discovery of sequences targeting the recently emerged variants (Delta and Omicron), while still maintaining their capability to bind and act upon the original variant as well as other coronavirus-related viral diseases (SARS-CoV, MERS-CoV and H1N1). Given the promising state of RNA technologies, exploring siRNAs as therapeutics would open the door to easily accessed personalized medicine, even to the individual level.

Also, despite current focus on nanostructures, delivery peptides are a promising delivery system, albeit the few number of described delivery peptides have restricted their use to the most common CPPs. Here, we have resorted to the power of deep learning to design new potential cell delivery peptides and explored the physicochemical properties to assign their activity prediction. The emergence of *in silico* tools and servers for these analyses are the way to go to enthusiasm researchers to better select and design peptide and nucleic acids sequences that could have better results in wet lab experiments, while helping reduce the cost of those tests via previous screening and selection. Thus, we designed 62 siRNA sequences targeting the most important SARS-CoV-2 genes, mainly to pp1ab and spike proteins, which are proteins with vital roles in the replication and host invasion, respectively. Additionally, we came up with 10 new putative delivery sequence systems that could be able to cross the cellular membrane. Some of these sequences displayed better features than the TAT control peptide, especially regarding the maximum recommended dose, which could point to a better safety and lower toxicity when compared to this control. The emergence of new variants and diseases may push the need to new therapies, but RNA and data analysis may lead the way to answer these needs.

## Acknowledgements

The authors acknowledge computing resources made available by the Minerva HPC from the Coimbra Institute of Engineering (ISEC); the National Distributed Computing Infrastructure (INCD) funded by Fundação para a Ciência e a Tecnologia (FCT) and European Regional Development Fund (ERDF) [01/SAICT/2016 n°. 022153]; and the Advanced Computing Project [CPCA/A2/4568/2020]. We would also like to acknowledge and thank Bruna Mendes for the revision of the methodology used in this work.

## Funding

This work was supported by the Portuguese National funds via Fundação para a Ciência e a Tecnologia (FCT) and Regional Operational Program Centro (CENTRO2020) under the Portuguese Partnership Agreement 2020 by the European Regional Development Fund (ERDF) [2020.10114.BD, SFRH/BD/144303/2019, IF/01272/2015, and UIDB/04539/2020].

## Data availability

The data underlying this article was accessed from public databases. Genomes were obtained from NCBI, ENSEMBL and GISAID (https://gisaid.org). Peptide’s data was obtained from CPPsite2.0 (http://crdd.osdd.net/raghava/cppsite), MLCPP (http://thegleelab.org/MLCPP), APD (http://aps.unmc.edu), CAMP (http://camp.bicnirrh.res.in), DBAASP (https://dbaasp.org), and LAMP (http://biotechlab.fudan.edu.cn/database/lamp). The derived data generated in this research will be shared on reasonable request to the corresponding author.

## Supporting Information

**Figure S1.**
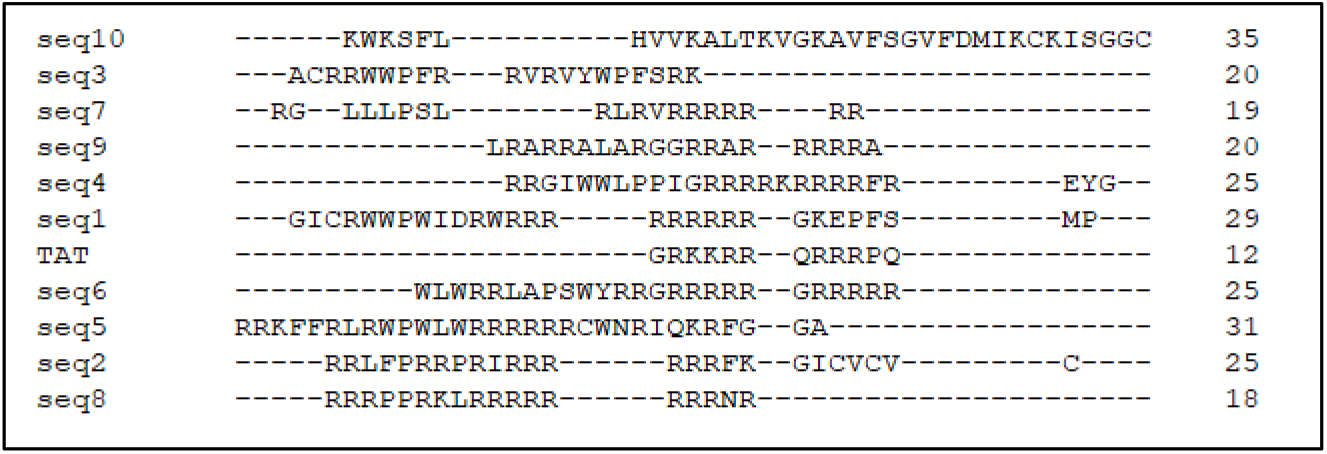
Sequence alignment of the new peptides sequence to TAT cell penetrating peptide.

**Table S1.** Distribution of annotated genes that the produced 21-nt sequences can target. Pp, polyprotein; nsp, non-structural protein; 3CL-PRO, 3C-like protease (main protease); ORF, open reading frame; ns, accessory protein.

| Gene/set of genes | Number of targets | Gene/set of genes | Number of targets |
| --- | --- | --- | --- |
| pp1ab, pp1a, nsp3 | 5811 | Envelope protein | 249 |
| pp1ab, polymerase | 2732 | Nucleocapside, ORF14 | 243 |
| Spike protein S1 | 2019 | pp1ab, pp1a, nsp7 | 225 |
| pp1ab, pp1a, nsp2 | 1890 | ns6 | 196 |
| pp1ab, helicase | 1779 | ns7b | 128 |
| pp1ab, exoribonuclease | 1557 | pp1ab, pp1a, polimerase, nsp11 | 39 |
| pp1ab, pp1a, nsp4 | 1476 | nsp1, pp1ab, pp1a, nsp2 | 24 |
| Spike protein S2 | 1751 | pp1ab, pp1a, nsp2, nsp3 | 24 |
| pp1ab, nsp15 | 1014 | pp1ab, pp1a, nsp4, 3CL-PRO | 24 |
| pp1ab, pp1a, 3CL-PRO | 894 | pp1ab, pp1a, 3CL-PRO, nsp6 | 24 |
| pp1ab, nsp16 | 894 | pp1ab, pp1a, nsp6, nsp7 | 24 |
| pp1ab, pp1a, nsp6 | 846 | pp1ab, pp1a, nsp8, nsp9 | 24 |
| ORF3a | 836 | pp1ab, pp1a, nsp9, nsp10 | 24 |
| Nucleocapside | 716 | pp1ab, pp1a, nsp10, polymerase, nsp11 | 24 |
| Membrane protein | 679 | pp1ab, polymerase, helicase | 24 |
| pp1ab, pp1a, nsp8 | 570 | pp1ab, helicase, exoribonuclease | 24 |
| nsp1, pp1ab, pp1a | 540 | pp1ab, exoribonuclease, nsp15 | 24 |
| pp1ab, pp1a, nsp10 | 393 | pp1ab, nsp15, nsp16 | 24 |
| ns8 | 335 | Spike protein S2, ORF3a | 13 |
| ORF7a | 317 | Membrane protein, ns6 | 11 |
| pp1ab, pp1a, nsp9 | 315 | ns8, nucleocapside | 7 |
| Nucleocapside, ORF-9b | 315 |  |  |

**Table S2.**
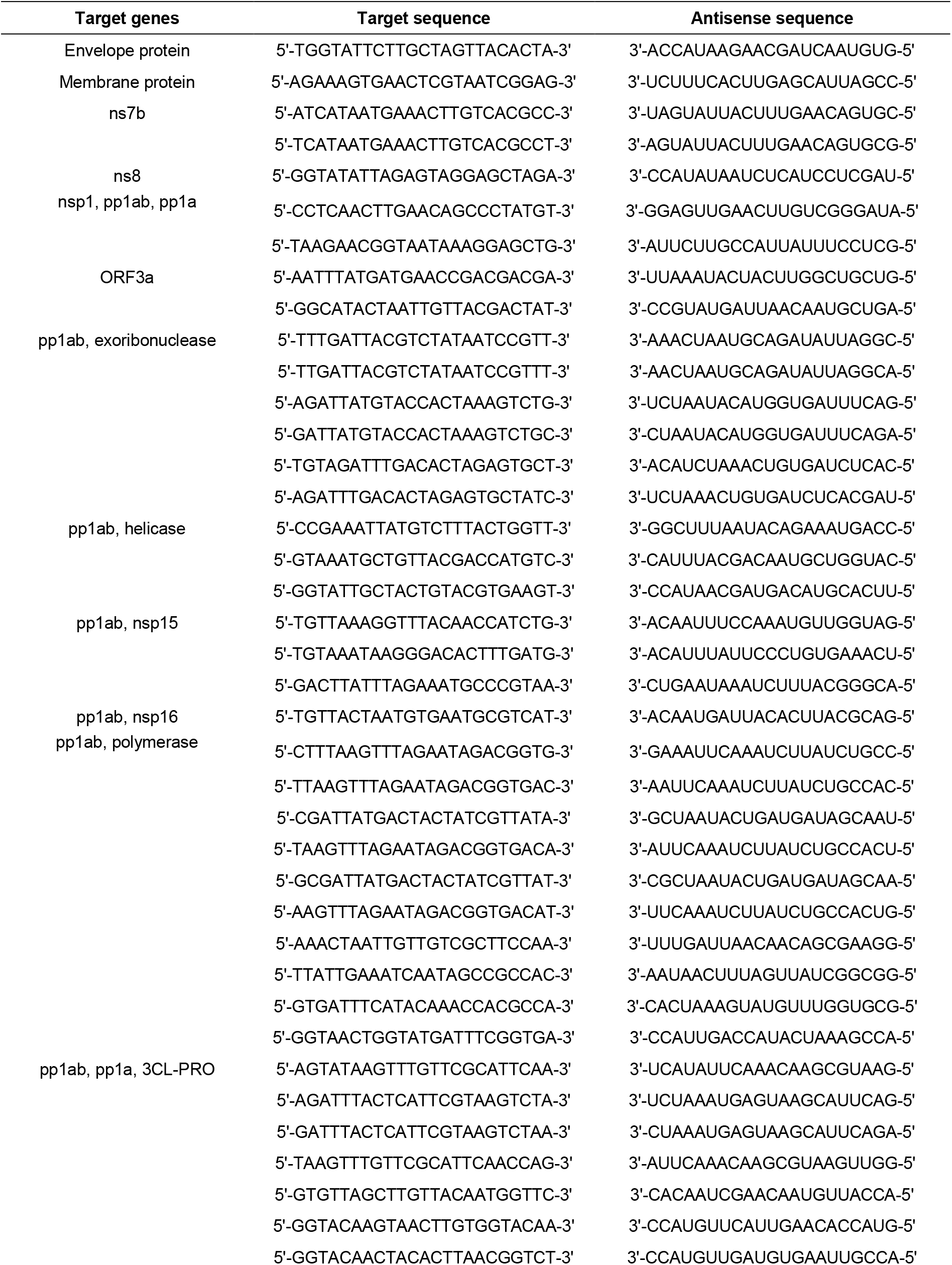
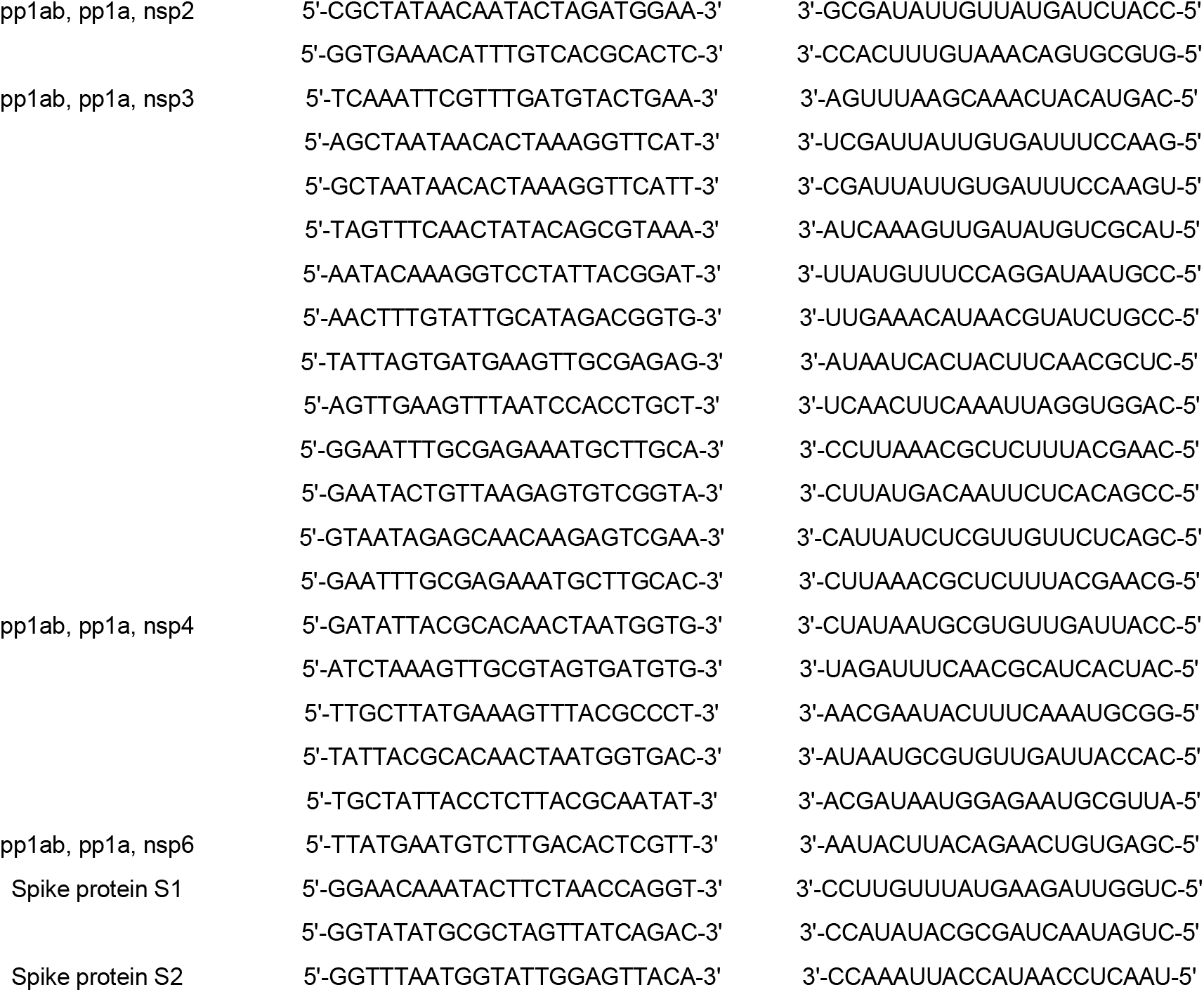
List of curated siRNAs targeting important viral genome genes. Pp, polyprotein; nsp, non-structural protein; 3CL-PRO, 3C-like protease (main protease); ORF, open reading frame; ns, accessory protein.

**Table S3.** Physicochemical features of the top 10 new CPP sequences, compared to the TAT CPP.

|  | Length<br>(aa) | Hydrophobicity | Hydrophobic<br>moment | Polar<br>residues (%) | Nonpolar<br>residues (%) | Net charge<br>(pH 7.4) |
| --- | --- | --- | --- | --- | --- | --- |
| **TAT** | 12 | -0.647 | 0.200 | 91.67 | 8.33 | 8 |
| **Seq1** | 29 | 0.199 | 0.180 | 58.62 | 41.38 | 10 |
| **Seq2** | 25 | 0.211 | 0.219 | 52.00 | 48.00 | 12 |
| **Seq3** | 20 | 0.497 | 0.058 | 50.00 | 60.00 | 7 |
| **Seq4** | 25 | 0.05 | 0.240 | 64.00 | 36.00 | 11 |
| **Seq5** | 31 | 0.233 | 0.217 | 58.06 | 41.94 | 14 |
| **Seq6** | 25 | -0.082 | 0.201 | 68.00 | 32.00 | 14 |
| **Seq7** | 19 | 0.016 | 0.024 | 63.16 | 36.84 | 10 |
| **Seq8** | 18 | -0.643 | 0.250 | 83.33 | 16.67 | 14 |
| **Seq9** | 20 | -0.308 | 0.235 | 65.00 | 35.00 | 11 |
| **Seq10** | 35 | 0.521 | 0.368 | 48.57 | 51.43 | 6 |

## Notes

### Competing Interest Statement

The authors have declared no competing interest.

## REFERENCES

(1) Piyush, R.; Rajarshi, K.; Chatterjee, A.; Khan, R.; Ray, S. Nucleic Acid-Based Therapy for Coronavirus Disease 2019. Heliyon 2020, 6 (9), e05007. https://doi.org/10.1016/j.heliyon.2020.e05007.

(2) Kumar, S.; Thambiraja, T. S.; Karuppanan, K.; Subramaniam, G. Omicron and Delta Variant of SARS-CoV-2: A Comparative Computational Study of Spike Protein. Journal of Medical Virology 2021, jmv.27526. https://doi.org/10.1002/jmv.27526.

(3) Hu, B.; Zhong, L.; Weng, Y.; Peng, L.; Huang, Y.; Zhao, Y.; Liang, X.-J. Therapeutic SiRNA: State of the Art. Sig Transduct Target Ther 2020, 5 (1), 101. https://doi.org/10.1038/s41392-020-0207-x.

(4) Houseley, J.; Tollervey, D. The Many Pathways of RNA Degradation. Cell 2009, 136 (4), 763–776. https://doi.org/10.1016/j.cell.2009.01.019.

(5) Torres-Vanegas, J. D.; Cruz, J. C.; Reyes, L. H. Delivery Systems for Nucleic Acids and Proteins: Barriers, Cell Capture Pathways and Nanocarriers. Pharmaceutics 2021, 13 (3), 428. https://doi.org/10.3390/pharmaceutics13030428.

(6) Taylor, R. E.; Zahid, M. Cell Penetrating Peptides, Novel Vectors for Gene Therapy. Pharmaceutics 2020, 12 (3), 225. https://doi.org/10.3390/pharmaceutics12030225.

(7) Nielsen, P. E.; Shiraishi, T. Peptide Nucleic Acid (PNA) Cell Penetrating Peptide (CPP) Conjugates as Carriers for Cellular Delivery of Antisense Oligomers. Artificial DNA: PNA & XNA 2011, 2 (3), 90–99. https://doi.org/10.4161/adna.18739.

(8) Duchardt, F.; Fotin-Mleczek, M.; Schwarz, H.; Fischer, R.; Brock, R. A Comprehensive Model for the Cellular Uptake of Cationic Cell-Penetrating Peptides. Traffic 2007, 8 (7), 848–866. https://doi.org/10.1111/j.1600-0854.2007.00572.x.

(9) Heitz, F.; Morris, M. C.; Divita, G. Twenty Years of Cell-Penetrating Peptides: From Molecular Mechanisms to Therapeutics: Peptide-Based Drug Delivery Technology. British Journal of Pharmacology 2009, 157 (2), 195–206. https://doi.org/10.1111/j.1476-5381.2009.00057.x.

(10) Ruseska, I.; Zimmer, A. Internalization Mechanisms of Cell-Penetrating Peptides. Beilstein J. Nanotechnol. 2020, 11, 101–123. https://doi.org/10.3762/bjnano.11.10.

(11) El-Andaloussi, S.; Järver, P.; Johansson, H. J.; Langel, Ü. Cargo-Dependent Cytotoxicity and Delivery Efficacy of Cell-Penetrating Peptides: A Comparative Study. Biochemical Journal 2007, 407 (2), 285–292. https://doi.org/10.1042/BJ20070507.

(12) Medeiros, I. G.; Khayat, A. S.; Stransky, B.; Santos, S.; Assumpção, P.; de Souza, J. E. S. A Small Interfering RNA (SiRNA) Database for SARS-CoV-2. Sci Rep 2021, 11 (1), 8849. https://doi.org/10.1038/s41598-021-88310-8.

(13) Fakhr, E.; Zare, F.; Teimoori-Toolabi, L. Precise and Efficient SiRNA Design: A Key Point in Competent Gene Silencing. Cancer Gene Ther 2016, 23 (4), 73–82.https://doi.org/10.1038/cgt.2016.4.

(14) Elbe, S.; Buckland-Merrett, G. Data, Disease and Diplomacy: GISAID’s Innovative Contribution to Global Health: Data, Disease and Diplomacy. Global Challenges 2017, 1 (1), 33–46. https://doi.org/10.1002/gch2.1018.

(15) Langmead, B.; Trapnell, C.; Pop, M.; Salzberg, S. L. Ultrafast and Memory-Efficient Alignment of Short DNA Sequences to the Human Genome. Genome Biol 2009, 10 (3), R25. https://doi.org/10.1186/gb-2009-10-3-r25.

(16) Tucs, A.; Tran, D. P.; Yumoto, A.; Ito, Y.; Uzawa, T.; Tsuda, K. Generating Ampicillin-Level Antimicrobial Peptides with Activity-Aware Generative Adversarial Networks. ACS Omega 2020, 5 (36), 22847–22851. https://doi.org/10.1021/acsomega.0c02088.

(17) Agrawal, P.; Bhalla, S.; Usmani, S. S.; Singh, S.; Chaudhary, K.; Raghava, G. P. S.; Gautam, A. CPPsite 2.0: A Repository of Experimentally Validated Cell-Penetrating Peptides. Nucleic Acids Res 2016, 44 (D1), D1098–D1103. https://doi.org/10.1093/nar/gkv1266.

(18) Manavalan, B.; Subramaniyam, S.; Shin, T. H.; Kim, M. O.; Lee, G. Machine-Learning-Based Prediction of Cell-Penetrating Peptides and Their Uptake Efficiency with Improved Accuracy. J. Proteome Res. 2018, 17 (8), 2715–2726. https://doi.org/10.1021/acs.jproteome.8b00148.

(19) Schyman, P.; Liu, R.; Desai, V.; Wallqvist, A. VNN Web Server for ADMET Predictions. Front. Pharmacol. 2017, 8, 889. https://doi.org/10.3389/fphar.2017.00889.

(20) Devi, G. R. SiRNA-Based Approaches in Cancer Therapy. Cancer Gene Ther 2006, 13 (9), 819–829. https://doi.org/10.1038/sj.cgt.7700931.

(21) Chernikov, I. V.; Vlassov, V. V.; Chernolovskaya, E. L. Current Development of SiRNA Bioconjugates: From Research to the Clinic. Front. Pharmacol. 2019, 10, 444. https://doi.org/10.3389/fphar.2019.00444.

(22) Wu, A.; Peng, Y.; Huang, B.; Ding, X.; Wang, X.; Niu, P.; Meng, J.; Zhu, Z.; Zhang, Z.; Wang, J.; Sheng, J.; Quan, L.; Xia, Z.; Tan, W.; Cheng, G.; Jiang, T. Genome Composition and Divergence of the Novel Coronavirus (2019-NCoV) Originating in China. Cell Host & Microbe 2020, 27 (3), 325–328. https://doi.org/10.1016/j.chom.2020.02.001.

(23) Ulferts, R.; Imbert, I.; Canard, B.; Ziebuhr, J. Expression and Functions of SARS Coronavirus Replicative Proteins. In Molecular Biology of the SARS-Coronavirus; Lal, S. K., Ed.; Springer Berlin Heidelberg: Berlin, Heidelberg, 2010; pp 75–98. https://doi.org/10.1007/978-3-642-03683-5_6.

(24) Samavati, L.; Uhal, B. D. ACE2, Much More Than Just a Receptor for SARS-COV-2. Front. Cell. Infect. Microbiol. 2020, 10, 317. https://doi.org/10.3389/fcimb.2020.00317.

(25) Safari, F.; Rahmani Barouji, S.; Tamaddon, A. M. Strategies for Improving SiRNA-Induced Gene Silencing Efficiency. Adv Pharm Bull 2017, 7 (4), 603–609. https://doi.org/10.15171/apb.2017.072.

(26) Elbashir, S. M. Functional Anatomy of SiRNAs for Mediating Efficient RNAi in Drosophila Melanogaster Embryo Lysate. The EMBO Journal 2001, 20 (23), 6877–6888. https://doi.org/10.1093/emboj/20.23.6877.

(27) Fedorov, Y.; Anderson, E. M.; Birmingham, A.; Reynolds, A.; Karpilow, J.; Robinson, K.; Leake, D.; Marshall, W. S.; Khvorova, A. Off-Target Effects by SiRNA Can Induce Toxic Phenotype. RNA 2006, 12 (7), 1188–1196. https://doi.org/10.1261/rna.28106.

(28) Matveeva, O. V.; Kang, Y.; Spiridonov, A. N.; Sætrom, P.; Nemtsov, V. A.; Ogurtsov, A. Y.; Nechipurenko, Y. D.; Shabalina, S. A. Optimization of Duplex Stability and Terminal Asymmetry for ShRNA Design. PLoS ONE 2010, 5 (4), e10180. https://doi.org/10.1371/journal.pone.0010180.

(29) Sievers, F.; Wilm, A.; Dineen, D.; Gibson, T. J.; Karplus, K.; Li, W.; Lopez, R.; McWilliam, H.; Remmert, M.; Söding, J.; Thompson, J. D.; Higgins, D. G. Fast, Scalable Generation of High-quality Protein Multiple Sequence Alignments Using Clustal Omega. Mol Syst Biol 2011, 7 (1), 539. https://doi.org/10.1038/msb.2011.75.

(30) Gautier, R.; Douguet, D.; Antonny, B.; Drin, G. HELIQUEST: A Web Server to Screen Sequences with Specific-Helical Properties. Bioinformatics 2008, 24 (18), 2101–2102. https://doi.org/10.1093/bioinformatics/btn392.

(31) Lee, A. G. Lipid–Protein Interactions in Biological Membranes: A Structural Perspective. Biochimica et Biophysica Acta (BBA) - Biomembranes 2003, 1612 (1), 1–40. https://doi.org/10.1016/S0005-2736(03)00056-7.

(32) Zou, L.-L.; Ma, J.-L.; Wang, T.; Yang, T.-B.; Liu, C.-B. Cell-Penetrating Peptide-Mediated Therapeutic Molecule Delivery into the Central Nervous System. CN 2013, 11 (2), 197–208. https://doi.org/10.2174/1570159X11311020006.

(33) Li, Z.-Q.; Jiang, L.-L.; Zhao, D.-S.; Zhou, J.; Wang, L.-L.; Wu, Z.-T.; Zheng, X.; Shi, Z.-Q.; Li, P.; Li, H.-J. The Modulatory Role of CYP3A4 in Dictamnine-Induced Hepatotoxicity. Front. Pharmacol. 2018, 9, 1033. https://doi.org/10.3389/fphar.2018.01033.

